# Investigating the molecular basis of cleptobiosis in eusocial stingless bees (Apidae: Hymenoptera)

**DOI:** 10.1101/2024.01.12.575469

**Authors:** Paulo Cseri Ricardo, Natalia De Souza Araujo, Maria Cristina Arias

## Abstract

Cleptobiosis, the act of raiding other species to obtain food or resources, is widespread among animals but rarely obligatory. The eusocial stingless bee *Lestrimelitta limao* is one of the few species that depend entirely on cleptobiosis, raiding other stingless bee colonies for survival. To investigate the molecular basis and evolutionary adaptations associated with this specialized lifestyle, we compared the transcriptomes of foraging workers of *L. limao* and three non-robber stingless bees – *Nannotrigona testaceicornis*, *Scaptotrigona* aff. *depilis*, and *Tetragonisca angustula*.

Our analysis revealed that differentially expressed orthologs were predominantly downregulated in *L. limao* workers, suggesting reduced transcriptional activity during foraging in this species. These downregulated genes fall into three major functional categories potentially linked to cleptobiotic adaptations: (1) detoxification and chemoreception genes, including cytochrome P450s and odorant receptors, indicating decreased exposure to phytochemicals; (2) neuronal and synaptic genes, such as *para* and *Dys*, possibly reflecting neurophysiological modifications; and (3) mitochondrial and carbohydrate metabolism genes, suggesting lower energetic demands. These findings provide novel insights into the molecular mechanisms shaping cleptobiotic behavior in eusocial bees.

## 1. Introduction

Cleptobiosis is an ecological relationship in which an individual or a group of individuals from the same species loot food or other valuable resources from another (Breed et al., 2012). This behavior is widespread in vertebrates and invertebrates and can be facultative or obligatory, although obligate cleptobiosis is far less common (reviewed in Iyengar 2008). Often the term cleptobiosis is associated with species of social insects, especially ants and bees (Breed et al., 2012). It was first proposed as a classification for the ecological interactions of thief ants (Forel, 1901; Wheeler, 1910), where several cases of inter and intraspecific food theft are known (Hölldobler, 1986; Perfecto and Vandermeer, 1993; Yamaguchi, 1995; Richard et al., 2004; Ronque et al., 2018; Yamaguchi, 1995).

Among eusocial bees, facultative cleptobiosis is common in species from the family Apidae encompassing the tribes Meliponini (stingless bees) and Apini (honeybees), especially when species are facing food shortage (Bohart, 1970; Costa et al., 2018; Michener, 2007; Wcislo, 1987). However, obligate cleptobiosis has only been confirmed in two genera of stingless bees: *Lestrimelitta* and *Cleptotrigona*. Species of *Trichotrigona* are also likely to show obligate cleptobiosis based on their nest architecture, but no direct observations of this behavior have been recorded so far (Camargo and Pedro, 2007; Pedro and Cordeiro, 2015). *Lestrimelitta* is a Neotropical genus comprising 26 described species, all exhibiting obligatory cleptobiotic behavior (Engel et al., 2023). In contrast, the Afrotropical genus *Cleptotrigona* includes only one extant species widely distributed across sub-Saharan Africa (Eardley, 2004), *Cleptotrigona cubiceps* (Lepeco et al., 2024; Portugal-Araújo, 1958). *Cleptotrigona* belongs to the Afrotropical Meliponini clade, while *Lestrimelitta* belongs to the Neotropical Meliponini clade (Rasmussen and Cameron, 2009). These two clades diverged over 70 million years ago (Rasmussen and Cameron, 2009). Thus, obligatory cleptobiosis in bees arose independently from flower-collecting ancestors at least twice.

*Lestrimelitta* and *Cleptotrigona* species attack nests of other stingless bee species to rob larval food, pollen, honey, and cerumen (Nogueira-Neto, 1997; Portugal-Araújo, 1958; Sakagami et al., 1993; Wittmann et al., 1990). Accordingly, these species have morphological and behavioral adaptations likely driven by their cleptobiotic behavior. Maybe the most striking of these adaptations is the absence of a functional corbicula, a morphological feature common to all corbiculate bees (Eardley, 2004; Michener, 2007; Shôichi F Sakagami and Laroca, 1963; Wille, 1983). Indeed, the reduction or complete loss of pollen-collecting structures seems to be a convergent adaptation in parasitic bees, as it is also observed in cleptoparasitic (*e.g.,* Nomadinae bees brood parasites) and social parasitic species (*e.g.*, *Bombus Psithyrus* subgenus) (Michener, 2000). Queens of *Lestrimelitta* have a higher number of ovarioles per ovary (10-15) than most stingless bees (∼4) (Sakagami, 1982), enabling the production of more workers. Moreover, the robust mandibles of the *Lestrimelitta* workers (James et al., 2022; Nogueira-Neto, 1997) help them overcome the colony defenses of host species.

An interesting behavioral adaptation in *Lestrimelitta* and *Cleptotrigona* species is their foraging strategy, as they carry out organized robbing raids in which several individuals simultaneously attack colonies of other species; in *C. cubiceps* the number of individuals participating in these raids may reach a few dozen (Portugal-Araújo, 1958), while in *Lestrimelitta* it could exceed several hundred (Sakagami et al., 1993). Raiding recruitment in *Lestrimelitta* has been reported to involve the use of semiochemicals (Blum, 1966; Sakagami et al., 1993) that have toxin functions (James et al., 2022), and may also play a role in the host response to raids (James et al., 2022; von Zuben et al., 2016). Likewise, in an apparent strategy of chemical deception, it has been suggested that *Lestrimelitta* may share cuticular hydrocarbons with their preferred host species to better conceal their presence (Vázquez et al., 2022).

Among all robber bees, *Lestrimelitta limao* is the best-known and most studied species (Rech et al., 2013). They are named after the citral volatile compound produced and released by workers, a compound common to all *Lestrimelitta* species (Blum et al., 1970; Francke et al., 2000). Like other species of *Lestrimelitta*, *L. limao* is specialized in raiding stingless bee nests (Wittmann et al., 1990), although records of pillaging in honeybee colonies exist (Sakagami et al., 1993). There are records of *L. limao* raids in more than a dozen stingless bee species from different genera, including *Friesella*, *Frieseomelitta*, *Melipona*, *Nannotrigona*, *Plebeia*, *Nannotrigona*, *Scaptotrigona*, *Tetragonisca* and *Trigonisca* (reviewed in Grüter et al. 2016). Nevertheless, *L. limao* host preference is apparently opportunistic depending on the location and the composition of the local stingless bee fauna (Sakagami and Laroca, 1963). *L. limao* raid attacks are common enough for this species to be considered a relevant commercial threat to bee-raising activities (meliponiculture), as it frequently attacks nests of species of commercial interest, killing many individuals and potentially leading to colony destruction (Sakagami et al., 1993; Sakagami and Laroca, 1963). In natural environments, it has been hypothesized that this species may control the size of stingless bee populations (Nogueira-Neto, 1997). In turn, evidence suggests that cleptobiotic interactions between *L. limao* and other stingless bee species may have driven the evolution of a soldier sub-caste in the assaulted species, which would be specialized in colony defense (Grüter et al., 2017, 2012). The number and size of these defensive soldiers are affected by the colony exposure to cleptobionts (Segers et al., 2016), corroborating the hypothesis that cleptobiosis would represent a relevant selective pressure in driving the evolution of host species.

Despite the evolutionary, ecological, and economic relevance of cleptobiosis in stingless bees, the mechanisms involved in the foraging strategy of *L. limao* and how they diverge from other non-parasitic bees are poorly understood. Likewise, the adaptations at the molecular level related to the evolution and maintenance of obligatory cleptobiosis in eusocial bees are unknown. Herein, we investigated the broad molecular mechanisms involved in stingless bees cleptobiosis using comparative transcriptomics. We generated transcriptomic data from foragers of the robber bee *L. limao* and for three pollen-collecting (non-parasitic) stingless bees, *Nannotrigona testaceicornis*, *Scaptotrigona* aff. *depilis*, and *Tetragonisca angustula*. These data were comparatively analyzed to identify and describe the molecular pathways associated with the cleptobiotic strategy of this robber bee during foraging.

## 2. Methods

### 2.1. Sampling and RNA sequencing

Workers were collected while performing foraging activities between 11:00 am and 1:00 pm to control for clock gene differences. We used foragers for the transcriptome comparative analyses as they are responsible for collecting food resources. The workers of *L. limao* were collected from three natural colonies in the city of São Paulo (Brazil); these workers were sampled when returning to their nests. Foragers of *N. testaceicornis* and *S.* aff. *depilis* were obtained from three rational colonies housed at the Bee Laboratory at the Universidade de São Paulo. For these species, workers were marked at their emergency and returned to their nests. After 30 days, when workers of several stingless bees are known to perform forager tasks outside the colony (see Grüter 2020), the marked bees were collected. For *T. angustula*, we used RNA-Seq data from two samples generated in a previous study, each consisting of a pool of three adult foragers collected using a similar sampling approach (Araujo and Arias, 2021). To account for potential sequencing batch effects, we additionally sequenced one batch sample from three *T. angustula* workers collected from a nest at the Instituto de Biociências, Universidade de São Paulo. These workers were collected while returning to their colony from foraging trips. Total RNA was extracted from each bee whole body using the RNeasy® Mini Kit (Qiagen). RNA extractions from individuals of the same nest were then pooled (3 individuals/pool) to reduce individual variation expression within the colonies. These pools are hereafter referred to as samples. In summary, three samples (one per colony) of *L. limao*, *N. testaceicornis,* and *S. depilis*; and one sample of *T. angustula* were sequenced. Library construction and sequencing were performed by Macrogen (South Korea) using the Illumina® NovaSeq 6000 platform and Illumina® suggested protocols to obtain 100 bp paired-end reads.

### 2.2. Transcriptome assembly

Read qualities were evaluated with FastQC 0.11.5 (Andrews, 2018). Raw reads were then quality-trimmed using Trimmomatic 0.38 (Bolger et al., 2014) (options: SLIDINGWINDOW:4:30 LEADING:3 TRAILING:3 MINLEN:80 AVGQUAL:30 HEADCROP:14). *De novo* transcriptome assembly was performed independently for each species using Trinity 2.10.0 (Grabherr et al., 2013; Haas et al., 2013), following default parameters, except for the *in silico* read normalization option (set up to 20x maximum coverage). The transcripts resulting from these assemblies were then clustered into SuperTranscripts (Davidson et al., 2017) using the Trinity gene splice modeler script implemented in the trinityrnaseq toolbox (Grabherr et al., 2013; Haas et al., 2013). Downstream analyses were performed using the SuperTranscripts datasets to reduce misassembly biases associated with the *de novo* approach. Quality parameters from SuperTranscripts were analyzed with Transrate 1.0.3 (Smith-Unna et al., 2016) and BUSCO 5.2.2 (Simão et al., 2015).

### 2.3. Annotation and orthologs identification

SuperTranscripts were annotated with Annocript 2.0.1 (Musacchia et al., 2015) using the UniProt Reference Clusters (UniRef90) (Suzek et al., 2015) and UniProtKB/Swiss-Prot (Bairoch, 1996) databases from February 2021. SuperTranscripts identified as possible contaminants (*i.e*., those annotated to Acari, Alveolata, Archaea, Bacteria, Fungi, Rhizaria, Rhodophyta, Viridiplantae and Viruses) were removed from the final data set using custom scripts (https://github.com/PauloCseri/Annotation.git). To identify orthologs among the four species, we followed the methodology from Ricardo et al. (2024). First, amino acid sequences derived from SuperTranscripts open reading frames (ORFs) generated by Annocript were used to identify orthogroups among all species using Orthofinder 2.1.1 (Emms and Kelly, 2019, 2015). Subsequently, for each orthogroup inferred by Orthofinder, only a one-to-one ortholog sequence per species was selected based on the best BLASTp alignment (BLAST 2.9.0+) (Altschul, 1997; Camacho et al., 2009) using default parameters.

### 2.4. Differential expression analyses

Differential expression (DE) analyses were performed using the data set of one-to-one orthologs of each species. To estimate the expression levels, the paired reads were mapped to the species correspondent set of ortholog sequences using Bowtie 2.3.5.1 (Langmead and Salzberg, 2012) and further processed with RSEM 1.3.3 (Li and Dewey, 2011). The identification of differentially expressed orthologs between *L. limao* and all other stingless bee species was pair-wise performed using two methods, as suggested by Ricardo et al. (2024): (1) using the TMM normalized counts (Robinson and Oshlack, 2010) compared in edgeR 3.34.0 (Robinson et al., 2010); and (2) using the scale based normalized (SCBN) counts (Zhou et al., 2019) and NOISeq 2.36.0 (Tarazona et al., 2015) for comparing expression levels. The SCBN method is a count normalization strategy optimized for cross-species DE analyses (Zhou et al., 2019) implemented in the Bioconductor package SCBN 1.10.0 (Zhou, 2021). Only significant differentially expressed transcripts (at a cut-off of |Log2FC| ≥ 2, adjusted p-value < 0.05) identified using both methods were filtered as DE orthologs. Lastly, we compared all pairwise differential expression results to identify orthologs consistently differently expressed between *L. limao* and all pollen-collecting species.

### 2.5. GO functional analysis

Among the DE orthologs identified between *L. limao* and all the other species, we identified GO-enriched terms using the Bioconductor package TopGO 2.20.0 (Alexa and Rahnenfuhrer, 2020). Redundant enriched terms for Biological Process (BP) were summarized in a two-level hierarchical GO set using REVIGO web server (Supek et al., 2011) and these hierarchical sets were represented in a chart generated using the CirGO 2.0 software (Kuznetsova et al., 2019).

## 3. Results

### 3.1. Transcriptome assembly and annotation

The RNA sequencing generated 254,713,432 raw reads for *L. limao*, 261,049,812 for *N. testaceicornis*, and 259,553,460 for *S.* aff. *depilis*. The number of reads decreased to 182,096,230, 187,198,886, and 185,326.996, respectively, after cleaning, For *T. angustula*, 89,081,896 new raw reads were sequenced additionally to 118,318,606 cleaned reads from a previous study (Araujo and Arias, 2021), resulting in a final set of 182,707,804 cleaned reads. The main quality parameters associated with each transcriptome assembly are summarized in Table 1.

**Table 1.**
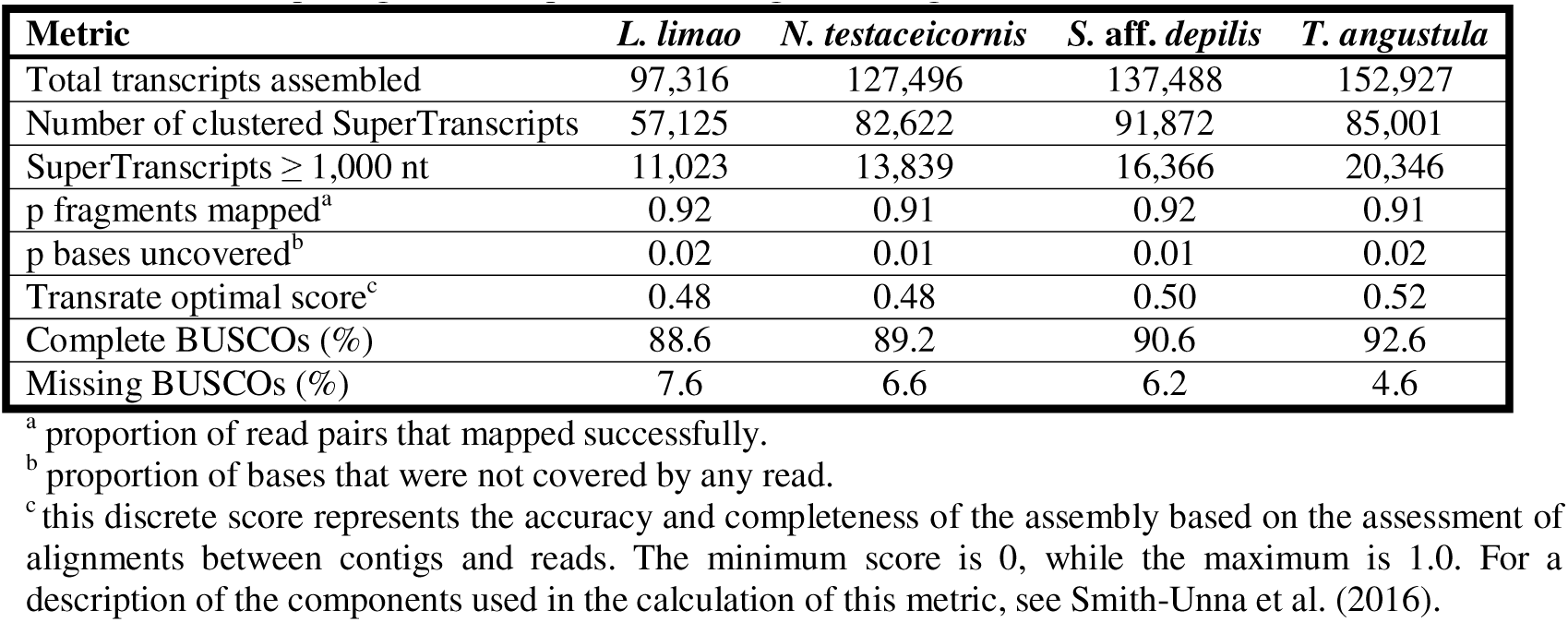
Main quality parameters for the transcriptome assemblies of Lestrimelitta limao, Nannotrigona testaceicornis, Scaptotrigona aff. depilis, and Tetragonisca angustula.

The number of transcripts assembled for *L. limao* was lower in comparison to the other species even though the amount of input RNA-seq reads was similar for all of them. Accordingly, after the annotation step and the exclusion of potential contaminants (Supplementary Material 1), *L. limao* was the species with the lowest number of SuperTranscripts returning significant blast hits with protein-coding gene sequences from the UniRef90 database, 24,859 (43,5% of total SuperTranscripts). For *N. testaceicornis*, *S.* aff. *depilis*, and *T. angustula* this numbers were 32,452 (39,2%), 38,273 (41,65%), and 33,596 (39,5%), respectively. Moreover, *L. limao* also presented a considerably lower number of potentially contaminating sequences (379) against *N. testaceicornis* (13,802), *S.* aff. *depilis* (14,414), and *T. angustula* (9,096). Most of the potential contaminants obtained for the non-cleptobiont bees, i.e., *N. testaceicornis*, *S.* aff. *depilis*, and *T. angustula*, resulted in significant blast hits against plant sequences (13,588, 14,033, and 8,467, respectively), particularly with plants of Fabaceae and Myrtaceae families, which is likely explained by their pollen-collecting behavior. *L. limao*, on the other hand, presented only 379 SuperTranscripts identified as potential contaminant sequences, of which 87 were from plants, and the remaining were attributed mainly to fungi and bacteria.

### 3.2. Differential expression of orthologs

Orthofinder identified 44,968 orthogroups, of which 8,213 were common to all four species (Figure 1A). *L. limao* showed the smallest number of species-specific orthogroups (140), and only 49 of them included at least one annotated transcript. We retrieved 7,482 one-to-one orthologs from all species. This set of orthologs was used for differential expression analyses. In the pair-wise comparisons, the most significant difference in expression was between *T. angustula* and *N. testaceicornis* (total of 1,229 DE orthologs), while the least amount of differentially expressed orthologs reported was between *S.* aff. *depilis* and *L. limao* (total of 734 DE orthologs). Finally, we identified 175 orthologs commonly differentially expressed between *L. limao* and all the other three bee species studied (Figure 1B, Supplementary Material 2), most of which (108) were consistently downregulated in *L. limao*. Among these, we find an odorant receptor (with significant BLAST hits to *Or1* from *Apis cerana* and LOC100741725 from *Bombus impatiens*), and several genes of the cytochrome P450 monooxygenase superfamily (*CYP* genes).

**Figure 1.**
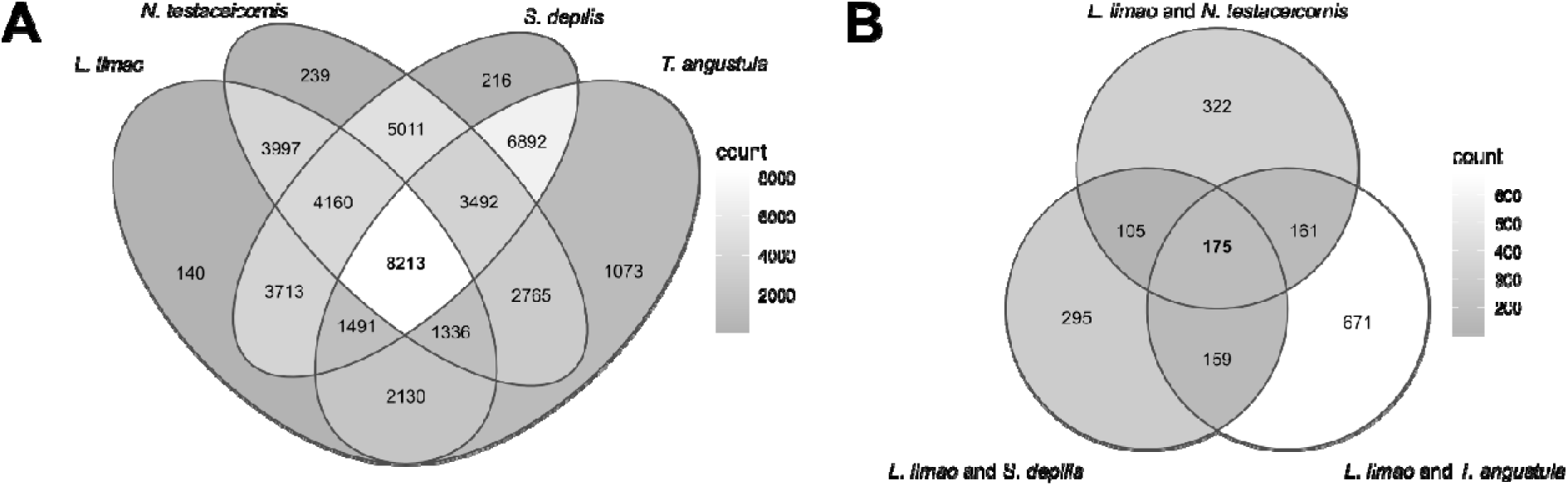
Analysis of the orthologs identified for the species *Lestrimelitta limao*, *Nannotrigona testaceicornis*, *Scaptotrigona* aff. *depilis* and *Tetragonisca angustula*. (A) Venn diagram representing the sets of orthogroups shared between/among species. (B) Venn diagram representing the sets of orthologs differentially expressed (DE) between/among species. Color scales represent count numbers ranging from lowest (dark gray) to highest (white).

The enrichment analysis of GO terms returned 26 enriched Biological Processes (BP) among the differentially expressed orthologs (Supplementary Material 3). The redundant BP terms were summarized in a two-level hierarchical list for visualization (Figure 2). The orthologs annotated to these enriched terms are shown in Table 2. Top enriched BP terms were related to *transmission of nerve impulse*, *maintenance of presynaptic active zone structure*, and *positive regulation of calcium ion transport*.

**Figure 2.**
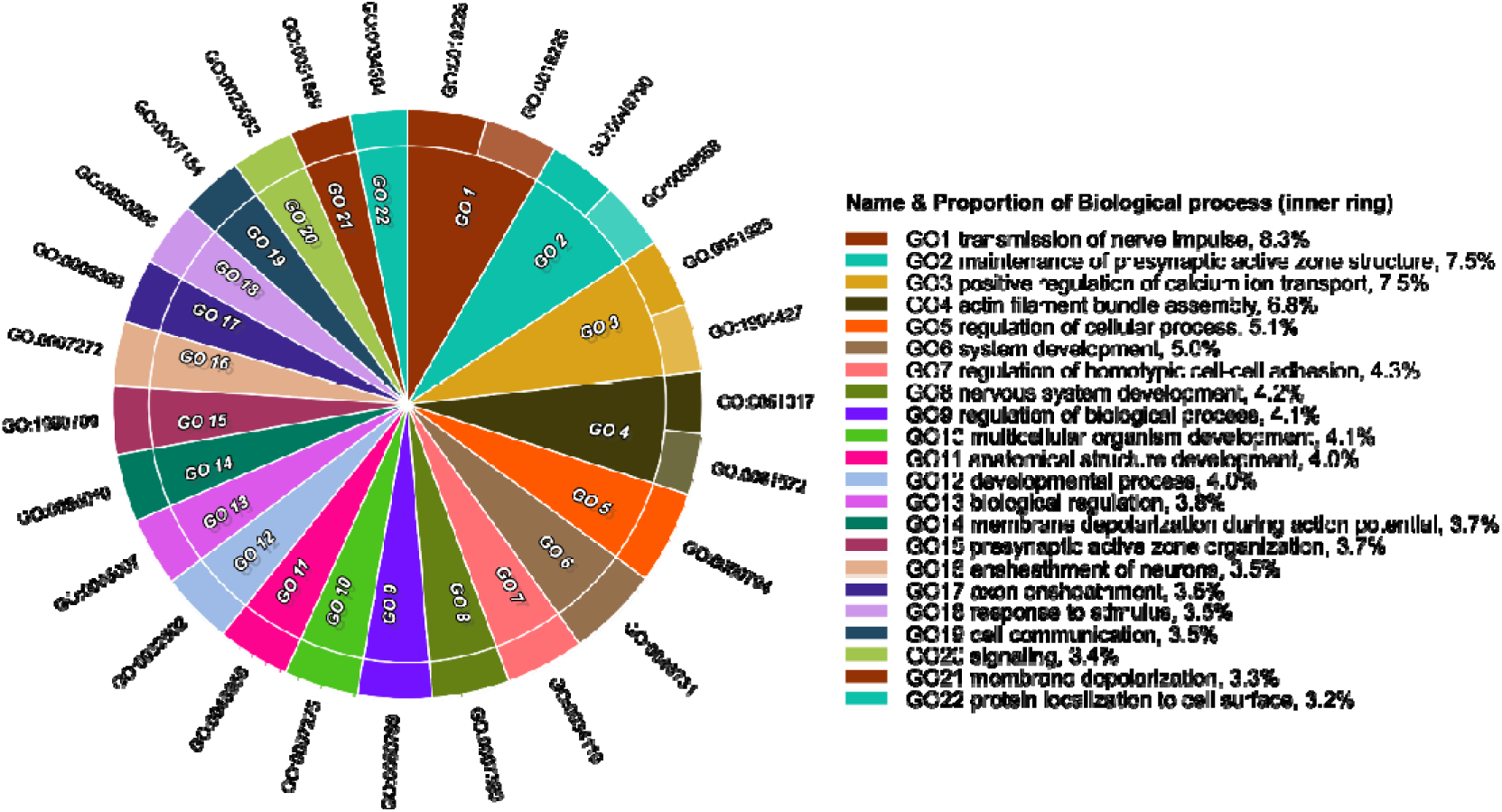
Summarized GO Biological Process terms enriched in the differentially expressed orthologs between the cleptobiotic *L. limao* and the pollen-collecting Meliponini species *Nannotrigona testaceicornis*, *Scaptotrigona* aff. *depilis* and *Tetragonisca angustula*. The legend next to the chart represents the color, name, and proportion of the GO parents for each two-level hierarchical set of child terms that were enriched.

**Table 2.**
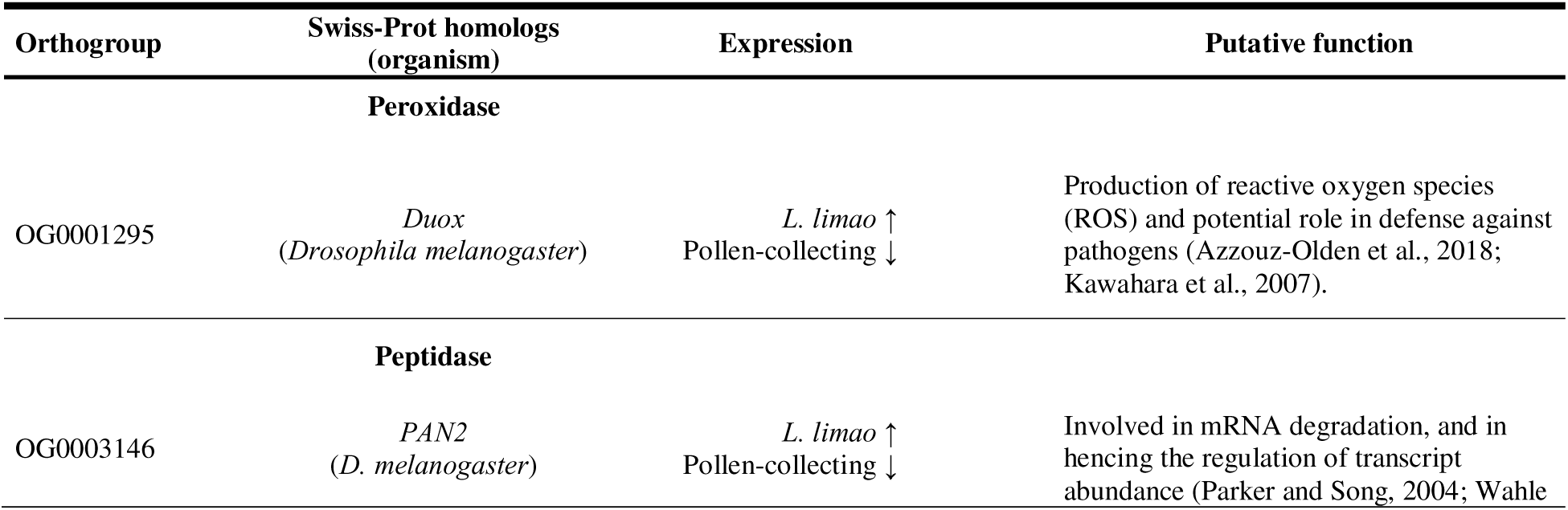
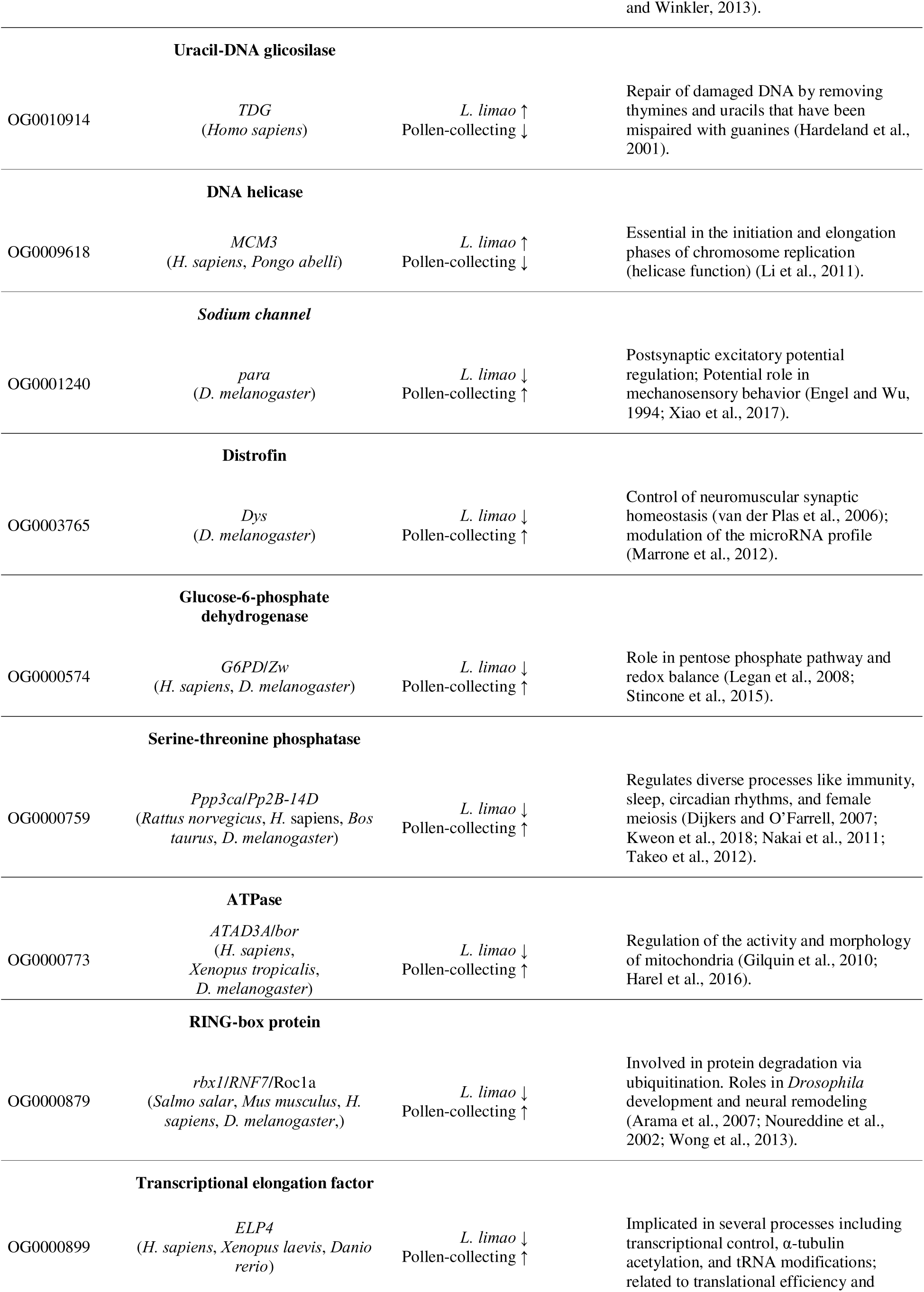
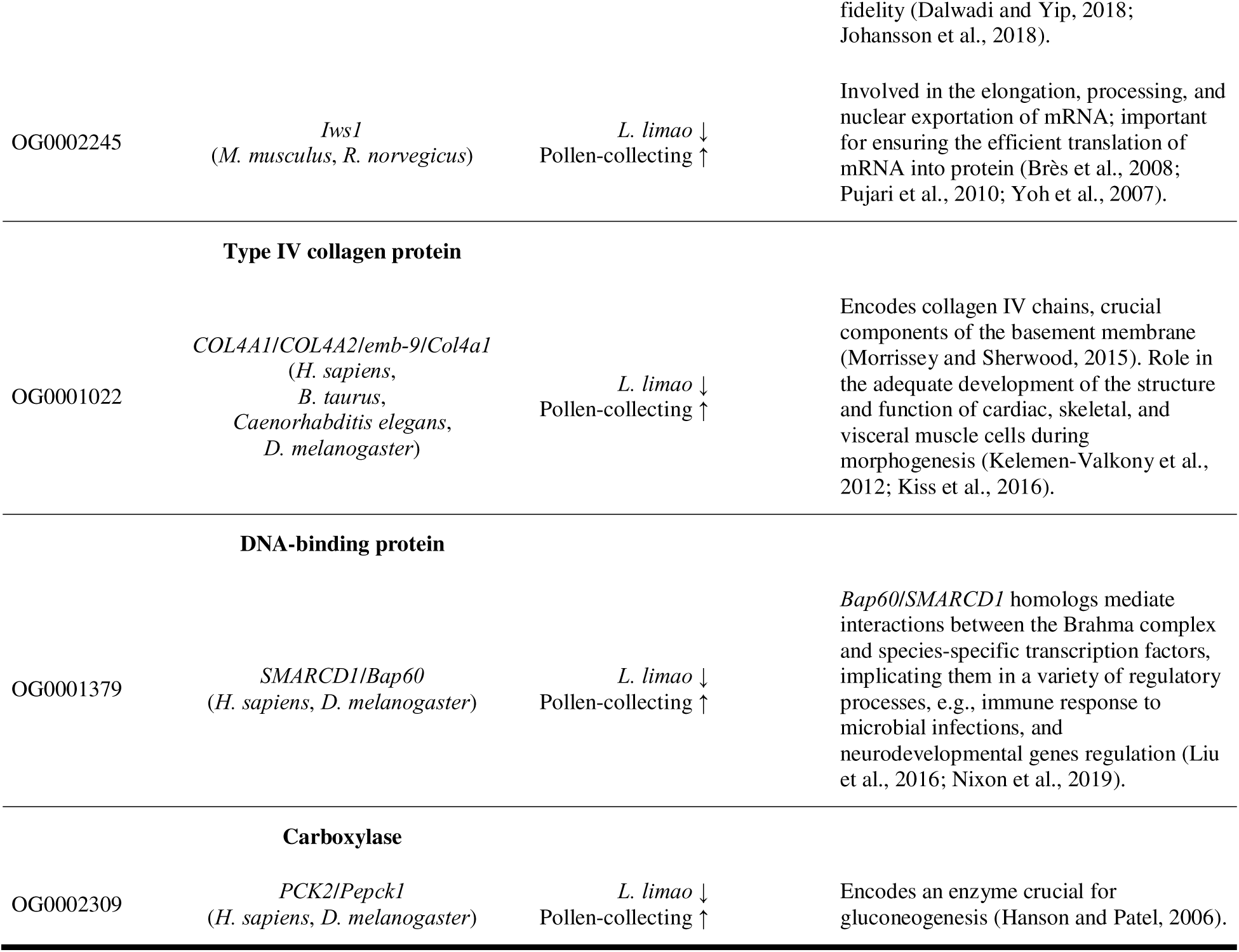
Differentially expressed orthologs between the cleptobiotic *L. limao* and the pollen-collecting Meliponini species *Nannotrigona testaceicornis*, *Scaptotrigona* aff. *depilis,* and *Tetragonisca angustula* annotated to the top enriched Biological Process GO term. Orthologs are represented by the ID of their respective orthogroups. Genes and respective species from the database hit are also shown (based on the Swiss-Prot database annotation) for each orthogroup. Expression patterns of orthologs are indicated by arrows: ↑ (upregulated) and ↓ (downregulated) as well as their function.

| Orthogroup | Swiss-Prot homologs (organism) | Expression | Putative function |
| --- | --- | --- | --- |
| **Peroxidase** |  |  |  |
| OG0001295 | *Duox*<br>(*Drosophila melanogaster*) | *L. limao* ↑<br>Pollen-collecting ↓ | Production of reactive oxygen species (ROS) and potential role in defense against pathogens (Azzouz-Olden et al., 2018; Kawahara et al., 2007). |
| **Peptidase** |  |  |  |
| OG0003146 | *PAN2*<br>(*D. melanogaster*) | *L. limao* ↑<br>Pollen-collecting ↓ | Involved in mRNA degradation, and in hencing the regulation of transcript abundance (Parker and Song, 2004; Wahle |
| **Uracil-DNA glycosylase** |  |  |  |
| OG0010914 | *TDG*<br>(*Homo sapiens*) | *L. limao* ↑<br>Pollen-collecting ↓ | Repair of damaged DNA by removing thymines and uracils that have been mispaired with guanines (Hardeland et al., 2001). |
| **DNA helicase** |  |  |  |
| OG0009618 | *MCM3*<br>(*H. sapiens*, *Pongo abelli*) | *L. limao* ↑<br>Pollen-collecting ↓ | Essential in the initiation and elongation phases of chromosome replication (helicase function) (Li et al., 2011). |
| **Sodium channel** |  |  |  |
| OG0001240 | *para*<br>(*D. melanogaster*) | *L. limao* ↓<br>Pollen-collecting ↑ | Postsynaptic excitatory potential regulation; Potential role in mechanosensory behavior (Engel and Wu, 1994; Xiao et al., 2017). |
| **Distrofin** |  |  |  |
| OG0003765 | *Dys*<br>(*D. melanogaster*) | *L. limao* ↓<br>Pollen-collecting ↑ | Control of neuromuscular synaptic homeostasis (van der Plas et al., 2006); modulation of the microRNA profile (Marrone et al., 2012). |
| **Glucose-6-phosphate dehydrogenase** |  |  |  |
| OG0000574 | *G6PD/Zw*<br>(*H. sapiens*, *D. melanogaster*) | *L. limao* ↓<br>Pollen-collecting ↑ | Role in pentose phosphate pathway and redox balance (Legan et al., 2008; Stincone et al., 2015). |
| **Serine-threonine phosphatase** |  |  |  |
| OG0000759 | *Ppp3ca/Pp2B-14D*<br>(*Rattus norvegicus*, *H. sapiens*, *Bos taurus*, *D. melanogaster*) | *L. limao* ↓<br>Pollen-collecting ↑ | Regulates diverse processes like immunity, sleep, circadian rhythms, and female meiosis (Dijkers and O'Farrell, 2007; Kweon et al., 2018; Nakai et al., 2011; Takeo et al., 2012). |
| **ATPase** |  |  |  |
| OG0000773 | *ATAD3A/bor*<br>(*H. sapiens*, *Xenopus tropicalis*, *D. melanogaster*) | *L. limao* ↓<br>Pollen-collecting ↑ | Regulation of the activity and morphology of mitochondria (Gilquin et al., 2010; Harel et al., 2016). |
| **RING-box protein** |  |  |  |
| OG0000879 | *rbx1/RNF7/Roc1a*<br>(*Salmo salar*, *Mus musculus*, *H. sapiens*, *D. melanogaster*,) | *L. limao* ↓<br>Pollen-collecting ↑ | Involved in protein degradation via ubiquitination. Roles in *Drosophila* development and neural remodeling (Arama et al., 2007; Nouredine et al., 2002; Wong et al., 2013). |
| **Transcriptional elongation factor** |  |  |  |
| OG0000899 | *ELP4*<br>(*H. sapiens*, *Xenopus laevis*, *Danio rerio*) | *L. limao* ↓<br>Pollen-collecting ↑ | Implicated in several processes including transcriptional control, $\alpha$ -tubulin acetylation, and tRNA modifications; related to translational efficiency and fidelity (Dalwadi and Yip, 2018; Johansson et al., 2018). |
| OG0002245 | *Iws1*<br>(*M. musculus*, *R. norvegicus*) | *L. limao* ↓<br>Pollen-collecting ↑ | Involved in the elongation, processing, and nuclear exportation of mRNA; important for ensuring the efficient translation of mRNA into protein (Brès et al., 2008; Pujari et al., 2010; Yoh et al., 2007). |
| **Type IV collagen protein** |  |  |  |
| OG0001022 | *COL4A1/COL4A2/emb-9/Col4a1*<br>(*H. sapiens*,<br>*B. taurus*,<br>*Caenorhabditis elegans*,<br>*D. melanogaster*) | *L. limao* ↓<br>Pollen-collecting ↑ | Encodes collagen IV chains, crucial components of the basement membrane (Morrissey and Sherwood, 2015). Role in the adequate development of the structure and function of cardiac, skeletal, and visceral muscle cells during morphogenesis (Kelemen-Valkony et al., 2012; Kiss et al., 2016). |
| **DNA-binding protein** |  |  |  |
| OG0001379 | *SMARCD1/Bap60*<br>(*H. sapiens*, *D. melanogaster*) | *L. limao* ↓<br>Pollen-collecting ↑ | *Bap60/SMARCD1* homologs mediate interactions between the Brahma complex and species-specific transcription factors, implicating them in a variety of regulatory processes, e.g., immune response to microbial infections, and neurodevelopmental genes regulation (Liu et al., 2016; Nixon et al., 2019). |
| **Carboxylase** |  |  |  |
| OG0002309 | *PCK2/Pepck1*<br>(*H. sapiens*, *D. melanogaster*) | *L. limao* ↓<br>Pollen-collecting ↑ | Encodes an enzyme crucial for gluconeogenesis (Hanson and Patel, 2006). |

## 4. Discussion

Our comparisons revealed that the cleptobiotic species exhibits lower expression levels for most differentially expressed orthologs. Among these downregulated orthologs, we highlight three groups associated with genes encoding for proteins that might play a role in the ecological differences among *L. limao* and non-robber bees. The first group comprises orthologs involved in detoxification and chemoreception pathways, including genes homologous to the cytochrome P450 monooxygenase superfamily and a putative odorant receptor (*Or1*). We hypothesize that a reduced expression of these genes could be associated with non-floral foraging lifestyle adaptations. The second group contains orthologs from the top enriched GO terms, *i.e.*, transmission of nerve impulses, maintenance of presynaptic active zone structure, and positive regulation of calcium ion transport. Although the exact functional consequences of these processes in cleptobiosis are unclear, the critical role of these genes in synaptic transmission, neuromuscular function, and sensory processing suggests they may be associated with neurophysiological adaptations of cleptoparasitism. Finally, the third group of orthologs is involved in mitochondrial activity and carbohydrate metabolism. We hypothesize that the reduced expression of these genes in *L. limao* workers suggests a shift in metabolic pathways leading to a low-energy-demanding worker state compared to pollen-collecting bees.

### 4.1. Group I of downregulated orthologs: detoxification and chemoreception

The cytochrome P450 superfamily (CYP genes) is widely conserved across living organisms (Werck-Reichhart and Feyereisen, 2000). Genes in this superfamily encode enzymes that catalyze a broad range of oxidative processes, including both endogenous and xenobiotic detoxification (Feyereisen, 2012). In arthropods, six major CYP clans have been identified, with the most prominent being the CYP2, CYP3, CYP4, and mitochondrial clans (Dermauw et al., 2020). Our results indicate that *L. limao* exhibits a reduced expression of orthologs from the CYP2 (*Cyp304a1*) and CYP3 (*Cyp6a14* and *Cyp6a2*) clans. The CYP2 clan is evolutionarily conserved and participates in key biological pathways, such as juvenile hormone and ecdysteroid biosynthesis (Bassett et al., 1997; Li et al., 2014; Namiki et al., 2005; Rewitz et al., 2006), as well as in the development of sensory structures (Willingham and Keil, 2004). The CYP3 clan is the most diverse among arthropods (Dermauw et al., 2020) and includes the CYP6 family, which comprises *Cyp6a14*, *Cyp6a2*, and several other genes involved in the metabolism of xenobiotics and phytochemicals (Feyereisen, 2012). Functional studies have demonstrated that *Cyp6a14* variants in *Tribolium castaneum* enhance detoxification capacity against plant-derived toxins (Ercan et al., 2020), while in *Aphis glycines*, increased expression of a *CYP6A14*-like gene confers resistance to pyrethroid insecticides (Xi et al., 2015). Similarly, *Cyp6a2* has been linked to resistance against organophosphates (Dunkov et al., 1997) and DDT (Pedra et al., 2004) in *Drosophila*.

In bees, Johnson et al. (2018) found evidence of positive selection in the genes of the CYP6AS subfamily (part of the CYP6A family, CYP3 clan). Based on these findings, the authors hypothesized that the role of these genes in metabolizing and detoxifying phytochemicals present in plant nectar, pollen, and resins (compounds that are toxic if not degraded) has enabled honeybees to consume a flavonoid-rich diet (Johnson et al., 2018; Liu et al., 2005; Mao et al., 2017). Since *L. limao* does not directly harvest floral resources but instead obtains them already processed through nest robbing (Michener, 1946; Shoichi F Sakagami and Laroca, 1963),— a behavior corroborated by the significantly lower number of plant-associated contaminant sequences in *L. limao* compared to pollen-collecting species in our dataset —we hypothesize that the downregulation of these CYP genes may reflect reduced exposure of this species to phytochemicals, as this behavioral shift likely decreases the need for flower detoxification mechanisms.

Another example of a downregulated ortholog in *L. limao*, potentially linked to the loss of floral foraging, comes from a putative odorant receptor (significant blast hits against *Or1* from *Apis cerana* and LOC100741725 from *Bombus impatiens*). *AcerOr1* has been implicated in detecting volatile floral scents (Zhao et al., 2014), and its downregulation agrees with a non-floral foraging lifestyle as *L. limao* workers likely do not rely on floral scents to obtain food. Still, the odorant receptor (OR) gene family is large and highly diverse in Hymenoptera (Obiero et al., 2021). Thus, it is worth mentioning that the identified orthologs may not have retained their functions across species. Chemosensory-related genes, including ORs, likely play crucial roles in intraspecific communication during nest-robbing events (Sakagami et al., 1993; von Zuben et al., 2016) and the specific cues guiding *L. limao* workers in robbing nests of diverse hosts remain unknown, as well as the impact of cleptobiotic behavior on the evolution of these gene families.

### 4.2. Group II of downregulated orthologs: synaptic transmission and neurosensory processing

Among the orthologs associated with the top enriched GO terms (*i.e.*, transmission of nerve impulses, maintenance of presynaptic active zone structure, and positive regulation of calcium ion transport) we found the genes *para* and *Dys*. In *Drosophila*, *para* encodes voltage-activated sodium channels (Loughney et al., 1989), which are essential for regulating postsynaptic excitatory potential (Xiao et al., 2017) and are widely expressed in the central nervous system of adult flies (Ravenscroft et al., 2020). Alternative splicing in this gene leads to channel variants of distinct properties (Lin et al., 2012, 2009; O’Dowd and Smith, 1996; Thackeray and Ganetzky, 1995), and mutations in *para* have been linked to seizure-like behavior, paralysis, and altered mechanosensory responses (Engel and Wu, 1994; Parker et al., 2011; Takai et al., 2020). The gene *Dys* encodes the protein Dystrophin, a conserved component of the dystrophin-glycoprotein complex (DGC) (Greener and Roberts, 2000; Neuman et al., 2005). In *Drosophila*, *Dys* is involved in neuromuscular synaptic homeostasis (Jantrapirom et al., 2019; van der Plas et al., 2006), photoreceptor axon migration (Marrone et al., 2011; Shcherbata et al., 2007; Zhan et al., 2010), and microRNA profile modulation (Marrone et al., 2012). The loss of functional *Dys* has been associated with age-dependent muscle degeneration and movement impairments (Shcherbata et al., 2007). The critical roles of *para* and *Dys* in synaptic transmission, neuromuscular function, and sensory processing suggest that these orthologs may contribute to neurophysiological adaptations associated with cleptoparasitism. However, their precise functional consequences remain unclear. Future studies integrating neurophysiological and behavioral analyses will be necessary to elucidate the impact of these transcriptional differences in robber bees.

### 4.3. Group III of downregulated orthologs: mitochondrial activity and carbohy-drate metabolism

Processes associated with mitochondrial activity and carbohydrate metabolism were also enriched in our GO term analysis and include downregulated orthologs in *L. limao* homologs to the vertebrate *ATAD3A*, to the human *PCK2,* and the *Drosophila bor*, *Pepck1*, *G6PD,* and *Zw*. *ATAD3A* plays a critical role in mitochondrial dynamics, particularly at the interface of the inner and outer mitochondrial membranes, where it regulates essential processes such as mitochondrial fusion and mitochondrial DNA organization (Gilquin et al., 2010; Harel et al., 2016; He et al., 2007; Hoffmann et al., 2009; Wang and Bogenhagen, 2006). Overexpression of *ATAD3A* enhances mitochondrial ATP production efficiency, a key driver of cellular metabolic activity (Gomes et al., 2011; Harel et al., 2016). *PCK2/Pepck1*, in turn, encodes an enzyme essential for gluconeogenesis (Hanson and Patel, 2006), playing a key role in glucose homeostasis in insects and initiating trehalose synthesis (Miyamoto and Amrein, 2017). Trehalose, the main sugar in insect hemolymph, is a crucial energy source for ATP production during flight (Becker et al., 1996; Matsuda et al., 2015). *G6PD* encodes glucose-6-phosphate 1-dehydrogenase, an enzyme critical for carbohydrate metabolism (reviewed in Stincone et al. 2015) and cellular redox balance (Kletzien et al., 1994; Legan et al., 2008). In this regard, it has been proposed that PPP may reduce the oxidative stress brought on by flying activity, particularly in species whose diets are rich in sugars, as is the case of insects that feed on nectar (Levin et al., 2017).

Given that these orthologs are linked to energic metabolism and protection against oxidative damage and that their expression levels are reduced in *L. limao* compared to non-robber bees, we conclude that the *L. limao* workers sampled in this study were not engaged in high-energy-demanding activities. We took special care to sample *L. limao* workers when returning to their colonies, this choice was based on the behavior of foragers from pollen-collecting species and ensured the sampling of workers at equivalent life stages across species for transcriptomic comparisons. However, *L. limao* workers are also known to raid other stingless bee colonies, and we did not sample individuals engaged in this activity. We hypothesize the expression pattern of workers may shift significantly during robbing raids since the metabolic demand should increase as worker activity intensifies. Accordingly, two interpretations are possible to explain our findings: either *L. limao* workers have a reduced global energetic demand compared to other stingless bees, or an alternative foraging state exists in this species. In this case, foraging workers would alternately engage in two distinct activities, robbing raids (a high-energy-demanding activity) and low-energy-demanding tasks, such as short-distance recognition flights. Further investigation comparing these two states is required to confirm these hypotheses.

## 5. Conclusions

*Lestrimelitta* species constitute one of the most notable and well-known examples of obligate cleptobiosis. Despite this, there is still a large gap in knowledge about the genetic factors underlying this behavior. Herein, we provide a starting point for these investigations. Our results demonstrated that most DE orthologs are downregulated in *L. limao* workers, and some of them may reflect specific aspects of obligate cleptobiosis in bees, such as the downregulation of *CYP* genes, which is associated with the non-floral foraging lifestyle. The enriched GO terms indicate that several DE orthologs potentially regulate neuronal and synaptic activities in addition to other important cellular processes. Moreover, we found evidence that robber foragers have reduced energetic demands when compared to other eusocial bee workers, which could suggest the existence of two foraging states in *L. limao,* one more active and another less active. In summary, the distinctive expression pattern of genes in *L. limao* provides valuable insights into the molecular mechanisms associated with its cleptobiotic behavior during foraging, offering avenues for further investigation into the adaptation and ecological dynamics of this unique species.

## Credit authorship contribution statement

**Paulo C. Ricardo:** Writing – review & editing, Writing – original draft, Conceptualization, Methodology, Sampling, Investigation, Formal analysis. **Natalia de S. Araujo:** Writing – review & editing, Conceptualization, Methodology, Supervision, Funding acquisition. **Maria C. Arias:** Writing – review & editing, Conceptualization, Supervision, Funding acquisition, Project administration.

## Acknowledgements

The authors acknowledge Susy Coelho Oliveira and Thiago Geronimo Pires Alegria for technical assistance, and Isabel Alves dos-Santos, Sheina Koffler, Larissa Nunes do Prado and Sergio Dias Hilário for support in specimen sampling.

## Data availability

The raw sequence reads have been deposited in the NCBI Sequence Read Archive (SRA) under the following accession numbers: SRR29109009, SRR29109008, SRR29109007, SRR29109006, SRR29109005, SRR29109004, SRR29109003, SRR29109002, SRR29109001, SRR29109000. The corresponding BioProject accession number is PRJNA1113344. The transcriptome assemblies, as well as the sets of SuperTranscripts, are available on FigShare (https://doi.org/10.6084/m9.figshare.26064586.v1).

## Declaration of generative AI and AI-assisted technologies in the writing process

During the preparation of this work the authors used ChatGPT (GPT-4o mini) and DeepSeek (DeepThink R1) in order to improve the clarity and coherence of the text and to assist in language editing and grammar correction. After using these tools, the authors reviewed and edited the content as needed, taking full responsibility for the content of the published article.

